# Ultra-high-speed multi-parametric photoacoustic microscopy

**DOI:** 10.1101/2021.12.25.474038

**Authors:** Fenghe Zhong, Song Hu

**Affiliations:** Department of Biomedical Engineering, Washington University in St. Louis; St. Louis, Missouri 63130, USA

## Abstract

Multi-parametric photoacoustic microscopy (PAM) is uniquely capable of simultaneous, high-resolution mapping of blood hemoglobin concentration, oxygenation, and flow *in vivo*. However, its speed has been limited by the dense sampling required for blood flow quantification. To overcome this limitation, we have developed an ultra-high-speed multi-parametric PAM system, which enables simultaneous acquisition of ~500 densely sampled B-scans by superposing the rapid laser scanning across the line-shaped focus of a cylindrically focused ultrasonic transducer over the conventional mechanical scan of the optical-acoustic dual foci. A novel optical-acoustic combiner is designed and implemented to accommodate the short working distance of the transducer, enabling convenient confocal alignment of the dual foci in the reflection mode. This new system enables continuous monitoring of microvascular hemoglobin concentration, blood oxygenation, and flow over a 4.5 × 3 mm^2^ area in the awake mouse brain with high spatial and temporal resolution (6.9 μm and 0.3 Hz, respectively).

## 1. Introduction

Capable of structural [1–3], functional [4], molecular [5], and metabolic imaging [6] with high spatial resolution *in vivo*, photoacoustic microscopy (PAM) is an emerging tool in biomedical research. Further, recent advances in multi-parametric acquisition and analysis make PAM uniquely capable of simultaneously mapping the total concentration of hemoglobin (C_Hb_), oxygen saturation of hemoglobin (sO_2_), and blood flow speed [6,7]. However, the dense sampling (*i.e.*, small scanning step size) required for blood flow quantification significantly limits the imaging speed of multi-parametric PAM, preventing dynamic assessments of microvascular function and tissue oxygen metabolism.

Currently, in multi-parametric PAM, the blood flow speed is quantified by analyzing the flow-induced decorrelation between adjacent A-line signals [8]. Although this method shows robust performance in measuring flow speeds across a wide range that is physiologically relevant (*i.e.*, 0.18—21 mm/s) and in blood vessels of different diameters [9], dense sampling (*i.e.*, 0.5 μm step size) is required to accurately extract the correlation decay constant. As a result, the B-scan rate is limited to ~1 mm/s, which prevents the use of existing high-speed acquisition methods developed for PAM [10–12] and sets up a barrier to improving the speed of multi-parametric PAM. Although other flow measurement methods have been developed or adopted for PAM, they have important limitations. For example, dynamically tracking the movement of individual blood cells is widely used for measuring microvascular flow [13,14]; however, this method is not readily applicable to flow quantification in large vessels where blood cells do not traverse in single files. Dual-pulse photoacoustic flowmetry based on the Grüneisen relaxation effect shows the promise of single-shot-based flow measurements [15]; however, an excessive average (*i.e.*, 100 times) is required, which significantly limits the imaging speed.

To improve the speed of multi-parametric PAM, we have adopted an optical-mechanical hybrid scan strategy to simultaneously acquire multiple B-scans while maintaining the dense sampling [16] for the correlation-based blood flow measurement. In our original demonstration, a spherically focused ultrasonic transducer was used [16]. Although the tight acoustic focus offers high sensitivity, the small acoustic detection area limits the range of laser scanning and thus the number of B-scans that can be simultaneously acquired. To overcome this limitation, we have replaced the spherically focused transducer with a cylindrically focused transducer [17]. However, the short working distance of the cylindrically focused transducer, which is required to achieve tight focus along one dimension for sufficient sensitivity, makes the integration of optical excitation and ultrasonic detection in the reflection mode a challenge. Although multiple different strategies have been developed for the integration of light and ultrasound in PAM, none of them can be readily applied to meet our development goal. For example, an acoustic reflector with a central opening for optical excitation was previously develop as an optical-acoustic combiner (OAC) for PAM [18], but that design is not compatible with the short working distance of the cylindrically focused transducer. The same difficulty is shared by other types of OACs that transmit light and reflect ultrasound, including an optically transparent acoustic reflector (*e.g.*, glass plate) [16], a dual-prism cube with a thin layer of silicone oil in between [19], and a single prism with an optical index-matching fluid [20].

Here, we report a new, small-footprint OAC and the enabled reflection-mode, ultra-high-speed, multi-parametric PAM system. Combining a high-repetition-rate pulsed laser, a high-speed resonant galvanometer, a cylindrically focused transducer, and the new OAC, the new multi-parametric PAM has achieved a 112-fold improvement in imaging speed over our previous system [16]—enabling simultaneous imaging of C_Hb_, sO_2_, and blood flow speed over an extended area of 4.5 × 3 mm^2^ in 3 s. The acquisition time can be further reduced to 0.5 s if the flow measurement (and thus dense sampling) is not required. The utility of this system has been demonstrated in the awake mouse brain by continuous monitoring of the dynamic changes of cerebrovascular blood oxygenation and flow in response to hypoxic challenges.

## 2. Methods

### 2.1 Optical-acoustic combiner

As shown in Fig. 1(a), the OAC is fabricated on a 100-μm acrylic film. A layer of aluminum (250 nm) is coated on the film to achieve light reflection in the visible spectral range. Then, a layer of SiO_2_ (100 nm) is coated on the top to protect the delicate aluminum coating, making it durable for experimental use in the water environment. In the PAM system (Fig. 1(b)), the OAC is mounted at 45° between the cylindrically focused transducer and the imaging target, with the use of a 3D-printed mount. The focused laser excitation beam is reflected by the aluminum layer onto the imaging target, while the generated ultrasonic wave back propagates through the OAC and is detected by the transducer. With a relatively low density (1.18 g/cm^3^) and acoustic velocity (2.8×10^3^ m/s), the acrylic has an acoustic impedance similar to the coupling medium (*i.e.*, water), which makes it an ideal substrate for the OAC to minimize the acoustic reflection. The thicknesses of the two coating layers are much smaller than the ultrasonic wavelength. Thus, they are also transparent to the light-generated ultrasonic wave.

**Figure 1.**
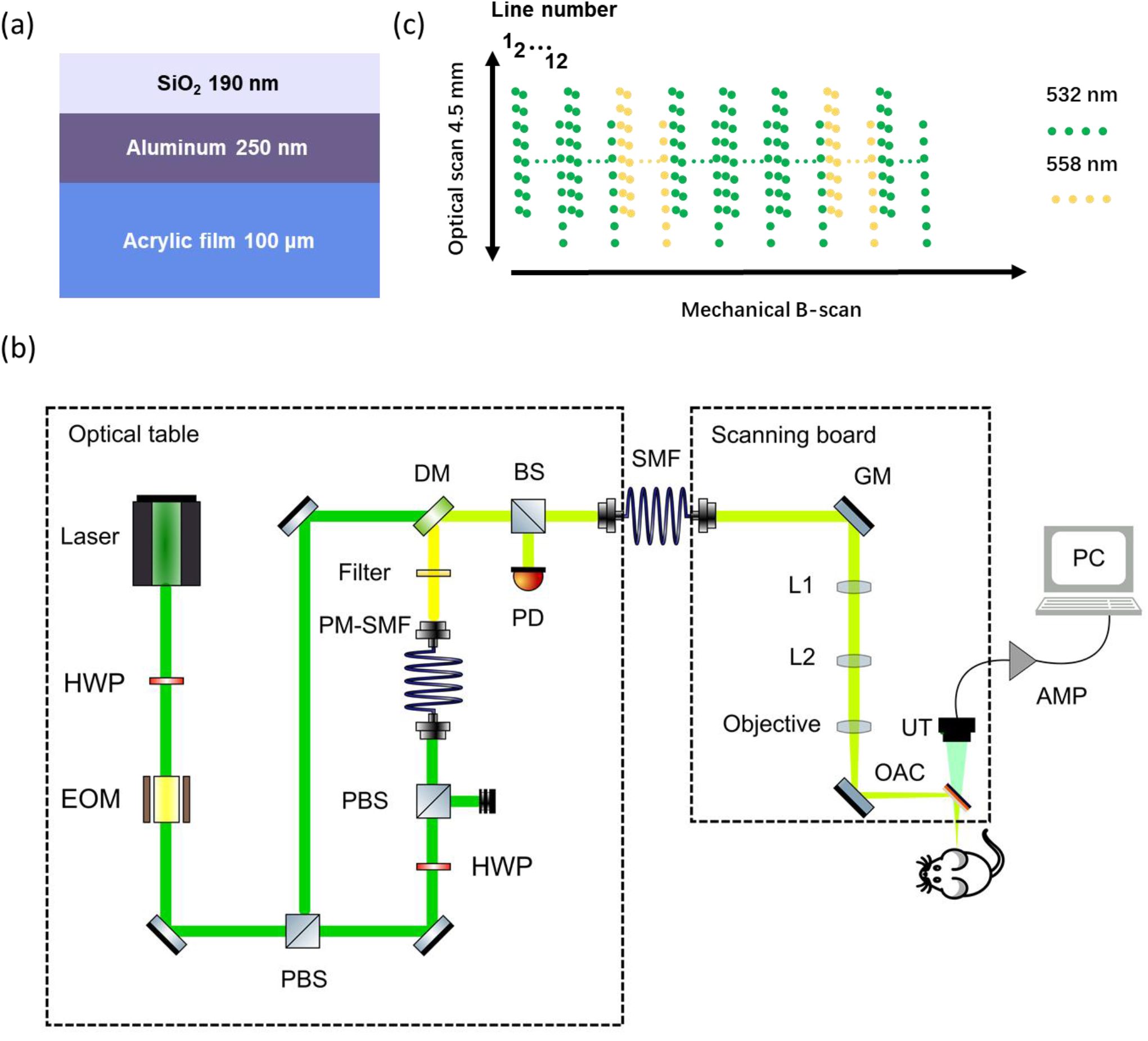
The new optical-acoustic combiner and enabled reflection-mode, ultra-high-speed, multi-parametric PAM. (a) Configuration of the OAC. (b) Scanning mechanism of the multi-parametric PAM. The EOM switches the wavelength from 532 nm to 558 nm after every 36 lines in the galvanometer (*i.e.*, optical) scanning direction. The green dots represent individual A-lines acquired at 532 nm, and the yellow dots represent those acquired at 558 nm. In the optical scanning direction, 83 A-lines are acquired in each round trip of the galvanometer scan. The mechanical scanning (*i.e.*, B-scan) direction is perpendicular to that of the optical scanning. (c) Configuration of the multi-parametric PAM. HWP, half-wave plate; EOM, electro-optic modulator; PBS, polarizing beam splitter; BS, beam sampler; PM-SMF, polarization-maintaining single-mode fiber; DM, dichroic mirror; PD, photodetector; GM, galvanometer; UT, ultrasound transducer; OAC, optical-acoustic combiner; AMP, amplifier.

### 2.2 Ultra-high-speed multi-parametric PAM

The configuration of the ultra-high-speed, multi-parametric PAM system is shown in Fig. 1(b). The pulses from a ns-pulsed laser (VGEN-G-20, Spectra-Physics; repetition rate, 1 MHz; wavelength, 532 nm) is dispatched to two different optical paths by an optical switch that consists of a half-wave plate (HWP; WPH05M-532, Thorlabs) and an electro-optical modulator (EOM; 350-80, Conoptics). When a low voltage is applied to the EOM, the light polarization remains unchanged. After passing through a polarizing beam splitter (PBS; PBS121, Thorlabs), the beam is coupled into a polarization-maintaining single-mode fiber (PM-SMF; HB450-SC, Fibercore; length, 8 m) through a fiber collimator (CFC-11X-A, Thorlabs) for wavelength conversion based on the stimulated Raman scattering effect. At the fiber output, a band-pass filter (BPF; ZET561/10X, Chroma) is applied to isolate the second Stokes peak at 558 nm. If a high voltage is applied to the EOM, the light polarization is rotated by 90°. After being reflected by the PBS, the 532-nm beam undergoes no wavelength conversion and is combined with the 558-nm Stokes beam via a dichroic mirror (DM; T550lpxr, Chroma). After 5% energy is tapped off by a beam sampler (BS; BSF10-A, Thorlabs) for the monitoring of laser fluctuation by a photodiode (PD; PDA25K2, Thorlabs), the dual-wavelength beam is coupled into the scanning head via a regular SMF (P1-460B-FC-1, Thorlabs; length, 1 m). In the scanning head, the dual-wavelength beam is collimated by a fiber collimator (CFC-11X-A, Thorlabs) and scanned by a resonant galvanometer (GM; 6SC12KA040-04Y, Cambridge). A lens pair (L1 and L2; AC254-050-A, Thorlabs) is used to relay the beam to the objective lens (AC127-050-A-ML). After being focused, the beam is reflected by a thin cut metallic mirror (PFR10-P01, Thorlabs) and the OAC onto the imaging target. The scanning direction of the galvanometer is carefully aligned to overlap the line focus of the transducer (UT; customized by the Ultrasonic Transducer Resource Center at the University of Southern California; Center frequency, 35 MHz) to maximize the sensitivity of ultrasonic detection.

### 2.3 Scanning strategy

The scanning strategy of the ultra-high-speed system is shown in Fig. 1(c). A resonant galvanometer steers the optical focus within the focus of the transducer at 12 kHz. When the laser runs at its maximum pulse repetition rate of 1 MHz, 83 A-lines are acquired across the 4.5-mm line focus in each round trip of the laser scanning. To ensure a reasonably small step size along this scanning direction, every 12 lines are grouped. In each group, the next line of sampling points is delayed with a constant time to generate a 10-μm displacement from the current line of points in the laser scanning direction. Thus, by combining the 12 lines, a step size of 10 μm can be achieved across the 4.5-mm laser scanning range. This strategy enables simultaneous acquisition of 498 B-scans along the mechanical scanning direction. Given the 12-kHz round-trip scanning rate and the grouping of 12 adjacent lines, the equivalent line rate is 2 kHz, which meets the sampling density required by the correlation-based blood flow measurement. By integrating the scanning mechanism and the cylindrically focused transducer, the imaging speed of multi-parametric PAM is improved by 112 times over our previous implementation with a spherically focused transducer [16].

For the sO_2_ measurement, the excitation wavelength is switched between 532 nm and 558 nm to distinguish the oxy- and deoxy-hemoglobin based on their optical absorption spectra. Specifically, after acquisition of 36 lines in the laser scanning direction, a high voltage is applied to the EOM to generate 558-nm light for acquisition of the next 12 lines. Two spatially adjacent A-lines acquired at the two different wavelengths are used for the sO_2_ calculation.

## 3. Results and discussion

### 3.1 Performance of the optical-acoustic combiner

The performance of the new OAC was evaluated by comprehensively quantifying its influence on the optical excitation and ultrasonic detection.

The optical reflectivity of the OAC at 532 nm and 558 nm were measured to be 85% with a power meter (PM100D, Thorlabs). To examine the influence of the OAC on the optical focus, we imaged the same resolution target (R1DS1P, Thorlabs) using the PAM system in both the reflection mode (with the OAC) and the transmission mode (without the OAC, as reported by us in [17]). As shown in Fig. 2(a), the lateral resolution is ~6.9 μm with or without the OAC, which indicates that the OAC has a good flatness or does not impair the beam quality. Together, these results suggest that the OAC has minimal influence on the optical excitation.

**Figure 2.**
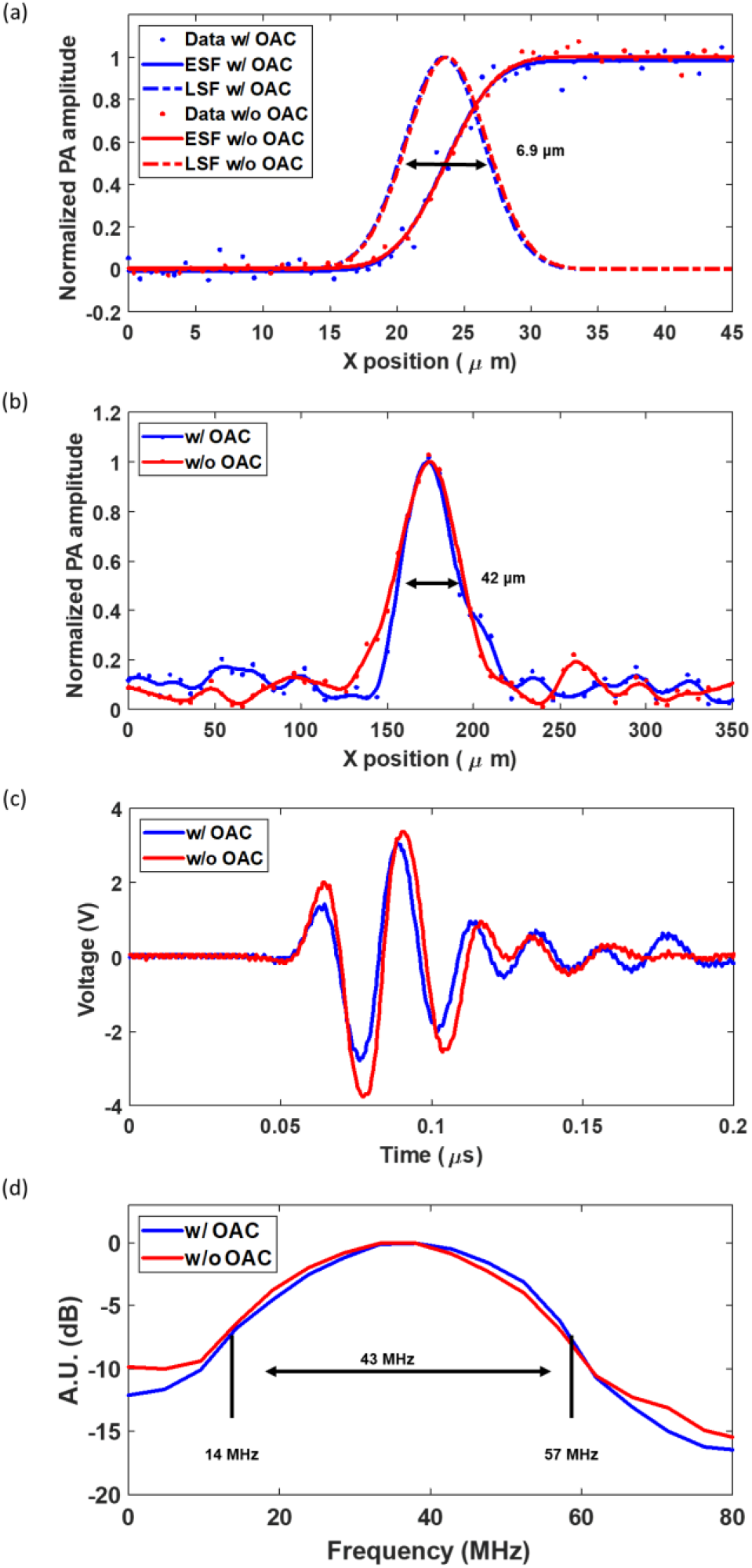
Influence of the OAC on the optical excitation and ultrasonic detection. (a) Lateral resolutions of the PAM with or without the OAC. (b) Axial resolutions of the PAM with or without the OAC. (c) Pulse-echo signals of the cylindrically focused transducer with or without the OAC. (d) Normalized frequency spectra of the pulse-echo signals with and without the OAC.

The axial resolution was quantified to be 42 μm with or without the OAC by analyzing the Hilbert transform of a representative A-line acquired from the resolution target (Fig. 2(b)). Since the axial resolution of PAM is determined by the ultrasonic bandwidth of the detected signal, this result suggests that the OAC does not affect the frequency spectrum of the photoacoustic signal. Moreover, a pulse-echo experiment was performed to quantify the influence of the OAC on the propagation of the ultrasonic wave. We used a pulser-receiver (5073PR, Olympus) to drive the cylindrically focused transducer. The generated ultrasonic wave was reflected by a glass slide and then received by the same transducer. Then, the measurement was repeated with the OAC inserted between the pulser-receiver and the glass slide. As shown in Fig. 2(c), the peak-to-peak amplitudes of the echo signals with and without the OAC are 5.9 V and 7.1 V, respectively, which indicates that the one-way ultrasonic attenuation of the OAC is < 9%. Further, Fourier analysis shows that the normalized frequency spectra of the echo signals with and without the OAC are nearly identical, both showing a 6-dB bandwidth of 43 MHz with the lower boundary at 14 MHz and higher boundary at 57 MHz (Fig. 2(d)). Together, these results suggest that the OAC has minimal influence on the ultrasonic detection.

### 3.2 *In vivo* imaging of the awake mouse brain

We tested the *in vivo* performance of the system in the brains of awake mice (CD-1, Charles River). A 3D-printed metal frame was installed on the mouse head after craniotomy by following our previously developed cranial window technique [21]. All animal procedures were carried out in conformity with the laboratory animal protocol approved by the Animal Care and Use Committee at Washington University in St. Louis.

First, the system was tested under the multi-parametric acquisition mode, where dense sampling was used for simultaneous acquisition of both sO_2_ and blood flow speed over an area of 4.5 × 3 mm^2^ at a frame rate of 0.3 Hz. After acquiring the baseline sO_2_ and blood flow under normoxia for 20 s, we switched the inhalation gas from medical air to hypoxic gas (10% oxygen), under which the mouse was continuously monitored for another 80 s. The high speed of the multi-parametric PAM clearly reveals the dynamic responses of the cerebral vasculature to the hypoxic challenge (Video 1). Side-by-side comparison of the two representative image sets acquired under normoxia and hypoxia (Figs. 3(a, c) and 3(b, d), respectively) shows that the hypoxic challenge resulted in a noticeable decrease in the sO_2_ and an attendant increase in the blood flow speed. These results echo our previous observations using a low-speed multi-parametric PAM system [6], which, however, could not provide the dynamic insights as shown in Fig. 3(e).

**Figure 3.**
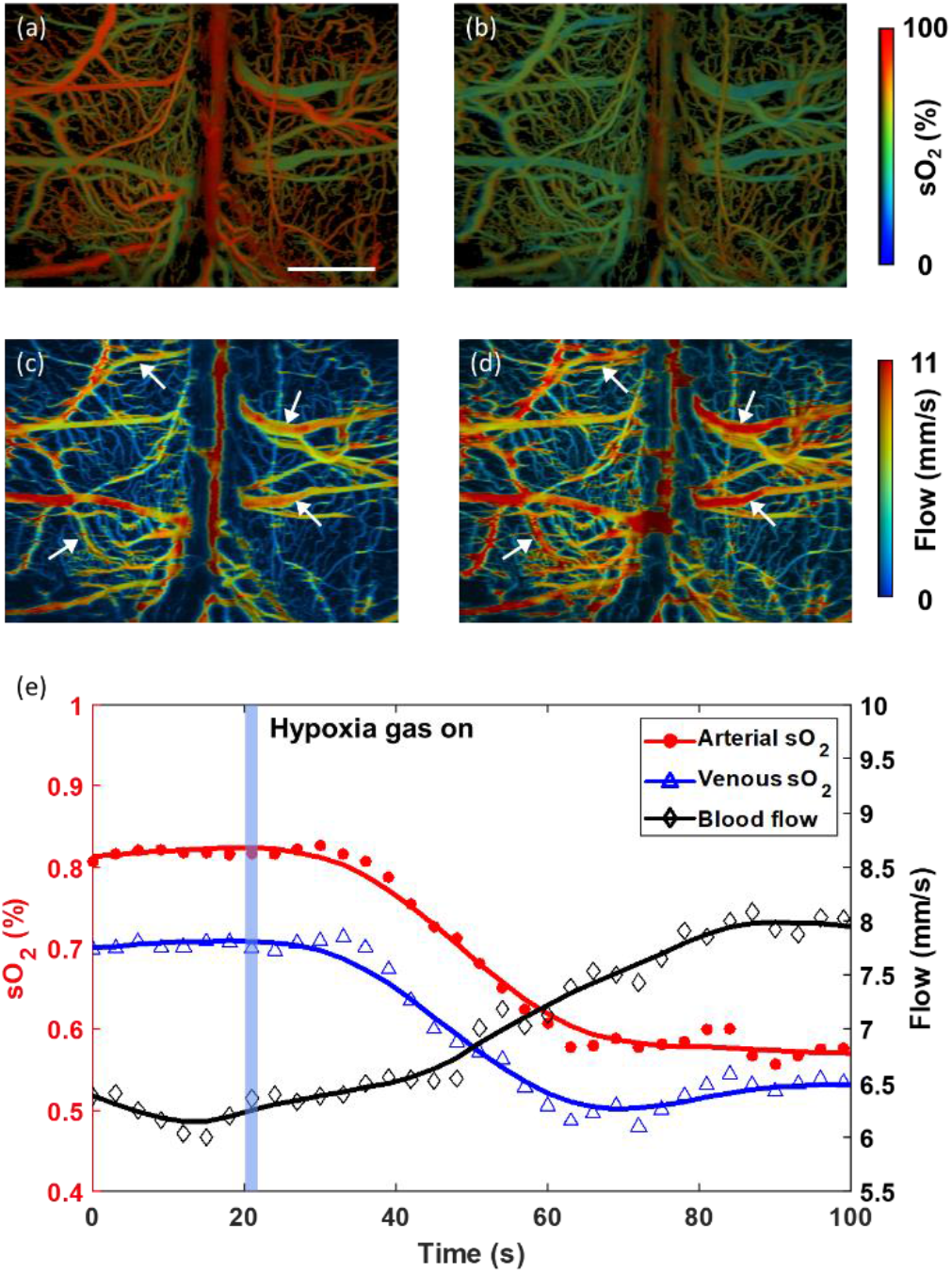
Ultra-high-speed, multi-parametric PAM of sO_2_ and blood flow speed over a 4.5 × 3 mm^2^ area in the awake mouse brain at a frame rate of 0.3 Hz. (a, b) sO_2_ images acquired under normoxia (left) and 100 s into hypoxia (right). (c, d) Blood flow images acquired under normoxia (left) and 100 s into hypoxia (right). (e) Dynamic responses of the average sO_2_ and blood flow to the hypoxic challenge. The red curve with dots and the blue curve with triangles shows the changes in arterial and venous sO_2_, respectively. The black curve with diamonds shows the change in blood flow speed. Scale bar: 1 mm.

In cases where blood flow measurement is not of interest, the speed of the system can be further increased by 6 times. To demonstrate this utility, the hypoxia experiment was repeated under the mode of sO_2_ acquisition only. As shown in Video 2, a cortical area of 4.5 × 3 mm^2^ was monitored at a frame rate of 2 Hz for 40 s, during which the inhalation gas was switched from medical air to hypoxic gas with 10% oxygen. Again, the representative sO_2_ images acquired under normoxia and hypoxia (Fig. 4(a) and 4(b), respectively) shows that the blood oxygenation decreases in response to the hypoxic challenge. Further, the high-speed monitoring clearly reveals the dynamic changes in the sO_2_.

**Figure 4.**
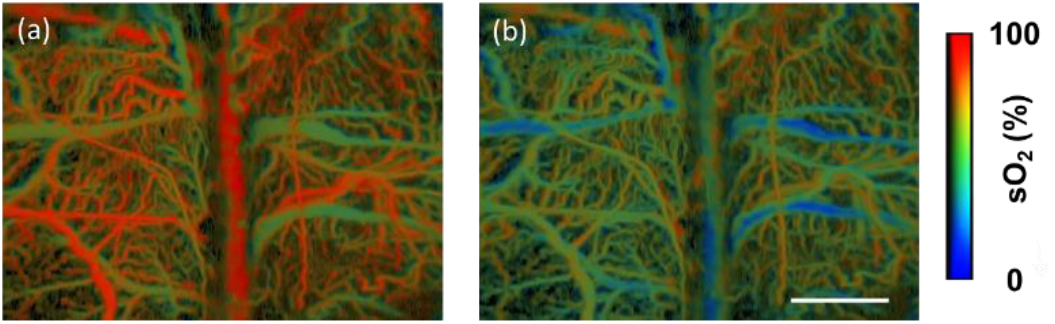
Ultra-high-speed, multi-parametric PAM of sO_2_ over a 4.5 × 3 mm^2^ area in the awake mouse brain at a frame rate of 2 Hz. (a) sO_2_ image acquired under normoxia. (b) sO_2_ image acquired 30 s into hypoxia. Scale bar: 1 mm.

## Conclusion

We have developed an ultra-high-speed multi-parametric PAM system with the unique ability to simultaneously image blood hemoglobin concentration, oxygenation, and flow at high spatial and temporal resolution over a large field of view. Our self-developed OAC with high optical reflectivity and acoustic transparency effectively accommodates the short working distance of the cylindrically focused transducer, thereby enabling the integration of the rapid laser scanning across the line-shaped acoustic focus and the conventional mechanical scan of the optical-acoustic dual foci in the reflection mode for simultaneous acquisition of 498 densely sampled B-scans. This system improves the speed of multi-parametric PAM by 112 times, enabling simultaneous mapping of blood oxygenation and flow over a 4.5 × 3 mm^2^ area in 3 s. We have demonstrated the utility of the system by monitoring the cerebrovascular responses to hypoxic challenges in awake mice. We believe that this new technique will find broad applications in basic and translational research.

## Supporting information

Video 1 Simultaneous monitoring of the blood oxygenation and flow responses of the awake mouse brain to hypoxic challenge. Scale bar: 1 mm

Video 2 Monitoring of the blood oxygenation response of the awake mouse brain to hypoxic challenge. Scale bar: 1 mm.

## Declaration of Competing Interest

The authors declare no conflicts of interest.

## Acknowledgements

The authors acknowledge the supports by the National Institutes of Health (NS099261 and NS120481), the National Science Foundation (2023988), and the Chan Zuckerberg Initiative DAF—an advised fund of Silicon Valley Community Foundation (2020-226174).

## Notes

### Competing Interest Statement

The authors have declared no competing interest.

